# *Bacillus subtilis* engineered for aerospace medicine: a platform for off-planet production of pharmaceutical peptides

**DOI:** 10.1101/2023.02.22.529550

**Authors:** Alec Vallota-Eastman, Cynthia Bui, Philip M. Williams, David L. Valentine, David Loftus, Lynn Rothschild

## Abstract

Biologics, such as pharmaceutical peptides, have notoriously short shelf lives, insufficient for long-duration space flight missions to the Moon or Mars. To enable the sustainable presence of humans on the Moon or Mars, we must develop methods for on-site production of pharmaceutical peptides in space, a concept we call *Astropharmacy*. Here, we present proof-of-concept for the first step needed: a low-mass system for pharmaceutical production designed to be stable in space. To demonstrate feasibility, we engineered strains of the space-hardy spore-forming bacterium, *Bacillus subtilis*, to secrete two pharmaceutical peptides important for astronaut health: teriparatide (an anabolic agent for combating osteoporosis) and filgrastim (an effective countermeasure for radiation-induced neutropenia). We found that the secretion peptides from the *walM* and *yoqH* genes of *B. subtilis* 168 worked well for secreting teriparatide and filgrastim, respectively. In consideration of the TRISH challenge to produce a dose equivalent in 24 hours, dried spores of our engineered strains were used to produce 1 dose equivalent of teriparatide from a 2 mL culture and 1 dose equivalent of filgrastim from 52 mL of culture in 24 hours. Further optimization of strain growth conditions, expression conditions, and promoter sequences should allow higher production rates to be achieved. These strains provide the template for future optimization efforts and address the first step in the *Astropharmacy*, capable of on-site production, purification, and processing of biopharmaceutical compounds in platforms amenable for use in space.

## 1 Introduction

As NASA prepares for long duration missions to the Moon and beyond, it must also prepare for the reality that astronauts will get sick hundreds of thousands of miles away from the nearest pharmacy. The Artemis Program—involving NASA, the European Space Agency, the Japanese Aerospace Exploration Agency and the Canadian Space Agency—seeks to re-establish a human presence on the Moon, and seeks to establish a basecamp to facilitate the human exploration of Mars and other deep space destinations (NASA, 2020; Batcha et al., n.d.). For short duration missions of up to a few months, or those where a quick return to Earth is possible, it is feasible for astronauts to take all the necessary medications they may need with them. The International Space Station (ISS) has ‘Med Kits’ for this reason, which include small supplies of painkillers, antibiotics, antiemetics, etc. (Medical Checklist, 2006). If needed, an ISS medical emergency can be dealt with by aborting the mission and having the crew return to Earth for recovery. However, the consideration of pharmaceutical availability becomes even more important for the multi-year missions that will be required for Mars, as speedy re-supply and/or evacuation missions are not possible. Reliance on pharmaceuticals is particularly important in space since invasive procedures, such as surgeries, are exceptionally challenging during flight (Blue et al., 2019). As such, it will be necessary to build the technological capability for on-site production of critical pharmaceuticals during the duration of the Artemis program and in preparation for human Mars missions.

Relying on pharmaceuticals launched with the crew is currently not possible for a planetary mission (Blue et al., 2019) for the following reasons. First, the shelf life of many pharmaceuticals is limited (Lyon et al., 2006). Of medications flown on ISS, 87% have shelf lives of fewer than 24 months. Shelf life in space may be even shorter because of degradation caused by exposure to the space environment (Kast et al., 2017). This problem becomes even greater when considering peptide pharmaceuticals, known as biologics. Biologics play a critical role in treating many of the medical conditions that astronauts are known to, or could likely, face. Unfortunately, biologics degrade in 6 months or less, even with refrigeration, further highlighting the problem of insufficient shelf-life for a Mars mission. A second problem is that, as the period of time away from Earth-bound pharmacies increases, the necessary range of medications of a rising variety of dosage and delivery forms increases, requiring greater up-mass. An ESA report (Berry et al., 2002) calculates the probability of a medical trauma is 0.06 per person per year. Our inability to store, transport and deliver pharmacologically-active biomolecules such as vaccines, antibodies, and other medications will limit long-term human missions. This has been recognized by NASA (NASA Technology Roadmaps - TA 6: Human Health, Life Support, and Habitation Systems, 2015). Additionally, NASA’s Human Research Roadmap identifies “Risk of Ineffective or Toxic Medications Due to Long Term Storage” as a major risk. The use of medicines in space flight also varies, since the way the body processes them is altered (Williams et al., 2003), as is their efficacy (Taylor, 2015). If we are to meet mission objectives of long-duration human missions including the exploration of Mars (Denis et al., 2020), and beyond, a new paradigm of pharmaceutical provision is required. A true paradigm shift would be to use life as a means of on-site production of any pharmaceutical an astronaut may need.

To fill this vital need, we have proposed the development of an “Astropharmacy”, a compact low-mass on-demand drug production system. Our concept consists of four parts: drug synthesis, purification, testing and administration. The first step will require a means of stable production, one that is low-mass, stable, and with reduced stringency for storage conditions. The second step will be to engineer a small and user-friendly device for purification of the drug, testing of the drug, and the final step will be to design a system for processing the drug to be ready for the required delivery form (e.g., intravenous, oral, dermal). The scope of this paper was to develop a proof of concept for the first step in this four-step process (cell-based biological production) while keeping in mind constraints imposed by the purification and processing steps that will be required in the future.

We chose *Bacillus subtilis* as the chassis organism for our proof of concept by engineering it to secrete filgrastim and teriparatide. While cell-free systems for peptide production were considered, a cell-based system was chosen at this point due to its intrinsic stability in space conditions (Zhang et al., 2020). *Bacillus subtilis* has long stood out as a promising chassis organism because *B. subtilis* endospores have been shown to survive the vacuum of space for nearly six years on NASA’s Long Duration Exposure Facility (LDEF) mission (Horneck, 1993b). The endospores are also able to survive long periods of desiccation, enabling storage of nearly massless dried spores (0.7 µg per million spores (Tisa et al., 1982; Cliff et al., 2005)) at room temperature. This “microbial astronaut” was also tested for its capabilities during the PowerCell payload experiment on Eu:CROPIS (Euglena and Combined Regenerative Organic-Food Production in Space) where it was successfully transformed with a selective plasmid in space (McCutcheon et al., 2016). In addition to its innate resistance to the hostile space environment, *B. subtilis* stands out in industrial settings for its ability to secrete various value-added products rather than retaining them inside the cell (van Dijl and Hecker, 2013; Liu et al., 2017; Gu et al., 2018; Zhang et al., 2020). Its robust secretory system means tht cell lysis is not necessary to harvest the product (Kakeshita et al., 2012), which makes purification easier. However, the relationship of secretion tag choice to the type of protein one would like to produce is still poorly understood. As a result, each peptide or protein one attempts to synthesize must be screened with all known signal peptides native to the *B. subtilis* genome.

Teriparatide and filgrastim are two important biologics, particularly for the perils of space flight (Fig. 1). These non-glycosylated peptides, stand out as the first therapeutic targets for on-site production due to their short shelf-life and easier route to synthesis. As proteins, they can be synthesized by cells such as bacteria. Further, native secretion mechanisms that cells use to transport proteins outside of the cytoplasm can be taken advantage of to produce extracellular biologics for ease of on-site separation from cell culture and purification.

**Figure 1.**
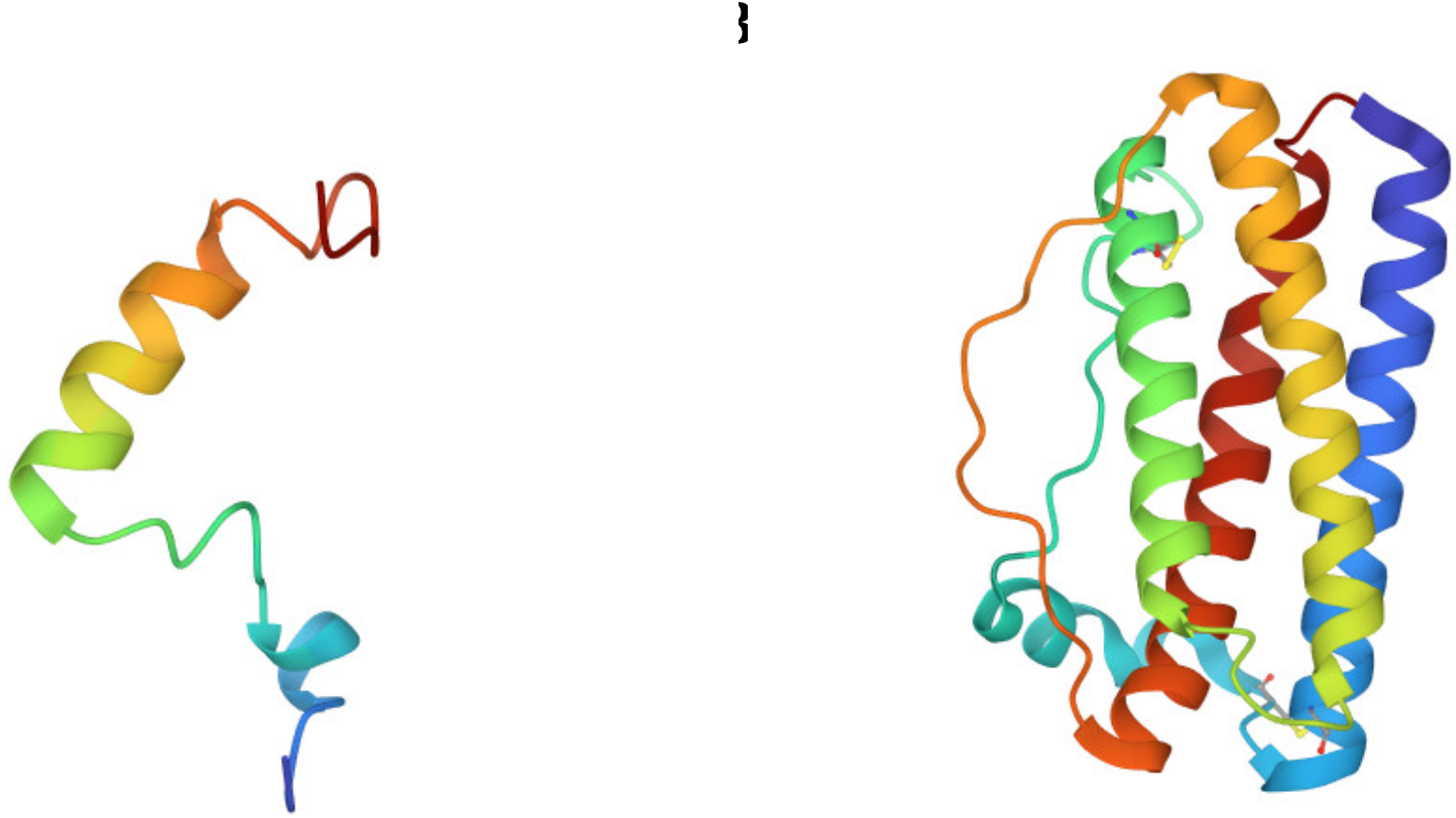
Ribbon diagram of teriparatide and filgrastim. A) Folded structure of 34 amino acid teriparatide. B) Folded structure of 175 amino acid filgrastim.

Teriparatide, also known as *recombinant human pyrathyroid hormone (1-34) (hPTH [1-34])* is the biologically active moiety of mineral homeostasis of the native full-length hormone hPTH (1-84) (Quattrocchi and Kourlas, 2004). This single chain polypeptide is a new FDA-approved biologic that increases bone mineralization and density. This agent could be used to counter the bone loss seen in space under reduced gravity conditions (Gabel et al., 2022). A single dose of teriparatide is 20 μg of the compound (Forteo, 2010).

Filgrastim, the name for human recombinant *granulocyte colony stimulating factor* (hG-CSF), is approved by the FDA and is now being used in the clinic to treat low-neutrophil blood count, a symptom of radiation toxicity. It is mostly used on Earth for improving radiation sickness in chemotherapy patients (Wright et al., 2017). Radiation exposure remains one of the most pressing issues in astrophysiology today (Chancellor et al., 2014). In the absence of Earth’s atmosphere, astronauts are exposed to a variety of radioactive particles, many of which cannot be effectively shielded by current shielding approaches. Astronauts on a 6 month-duration mission on ISS are exposed to approximately 75 millisieverts (mSv) of radiation – the equivalent radiation exposure of 375 chest x-rays (Hodkinson et al., 2017). Humans on Earth’s surface average 2.2 mSv of radiation per year (Hodkinson et al., 2017). By contrast, a 3-year mission to Mars, will expose astronauts to low-dose galactic cosmic rays, leading to radiation doses more than 13 times greater than astronauts experience on the ISS on average (Cucinotta and Durante, 2006; Afshinnekoo et al., 2020). This 1 Sv exposure is the equivalent of 5,000 chest x-rays. Notable known risks of excessive and/or prolonged exposure to radiation include cancer and radiation sickness. Filgrastim could be used to support the hematopoietic system by helping to restore neutrophil counts in the event of an acute radiation exposure from a solar particle event (especially if a crew member were to be un-shielded, on EVA). A single dose of G-CSF is 300 μg and it needs to be given once a day (Appelbaum, 1989).

The goal of this proof of concept was to screen a wide variety of *B. subtilis* secretion peptides for their ability to secrete filgrastim and teriparatide. We then modified the *B subtilis* genome to include the best-performing construct under a high constitutive expression promoter. Finally, as we were inspired by NASA’s Translational Research Institute for Space Health (TRISH) program (Mars, 2017), we set out to achieve TRISH’s goal of a production rate of at least one dose equivalent of each pharmaceutical peptide within 24 hours.

## Materials and Methods

### 3.1 Bacterial Growth

Bacterial strains used in all aspects of this project were *E. coli* HST08 Stellar™ Competent Cells and *B. subtilis* RIK1285 from TAKARA Bio. Growth for both *E. coli* and *B. subtilis* was carried out at 37°C in Luria Broth (LB) containing selective antibiotics with shaking at 200 rpm unless stated otherwise. For solid media, LB plates containing 1.5% agar were used. For pBE-S-based constructs (see Supplementary File 1 for list of abbreviations and meanings), ampicillin [100 µg/mL] was used for plasmid propagation in *E. coli* and kanamycin [10 µg/mL] was used for plasmid propagation in *B. subtilis*. For pECE321-based plasmid propagation, ampicillin [100 µg/mL] was used for plasmid propagation in *E. coli* and streptomycin [100 µg/mL] was used for selecting for genomic insertions in *B. subtilis*. To select for double-cross over events, erythromycin [1 µg/mL] was used as a counterselection by cross patching on solid media. For spore formation, Difco Sporulation Medium (DSM) was prepared according to the following recipe (Munakata et al., 1991). Per liter: Bacto Nutrient Broth (8g), 10% (w/v) KCl (10 mL), 1.2% (w/v) MgSO4×7H2O (10 mL), 1 M NaOH (~1.5 mL or to pH 7.6). Deionized (DI) water was added to 1 L and the solution was autoclaved, then allowed to cool to 50°C. Just prior to use, the following sterile solutions were added along with selective antibiotics: 1 M Ca(NO3)2 (1 mL), 0.01 M MnCl2 (1 mL), 1 mM FeSO4 (1 mL).

### 3.2 Cloning

PCR reactions were performed using PrimeSTAR® Max DNA Polymerase (TAKARA Bio Inc., San Jose, CA) according to manufacturer protocols. Homology cloning was performed using In-Fusion® Snap Assembly Master Mix for homology cloning according to manufacturer protocols (TAKARA Bio Inc., San Jose, CA). All PCR purifications were performed using the NucleoSpin® Gel and PCR Clean-up kit (TAKARA Bio Inc., San Jose, CA) according to manufacturer protocols. Plasmid preps from *E. coli* strains were performed using the PureYield™ Plasmid Miniprep System (Promega Corp., Madison, WI) according to manufacturer protocols. For plasmid preps from *B. subtilis* strains, water used to resuspend cells in the manufacturer’s protocol contained freshly dissolved lysozyme at 2.5 mg/mL final concentration and samples were incubated at 37 °C for 30 minutes before continuing with the normal protocol. DNA was quantified *via* the Thermo Scientific NanoDrop UV-Vis spectrophotometry system. Sanger sequencing was conducted by Elim Biopharmaceuticals, Inc. (Elim Biopharmaceuticals Inc., Hayward, CA).

#### 3.2.1 pAVE001 and pAVE003 Construction

Two expression plasmids were created for expression of the drugs: pAVE001 for filgrastim and pAVE003 for teriparatide, using basic homology-based cloning. gBlocks of drug cassettes were synthesized (Integrated DNA Technologies, Coralville, Iowa, USA). pBE-S (TAKARA Bio Inc.), the expression plasmid backbone, was linearized by PCR using primers F1/R1 (see Supplementary File 1 for primer table). For pAVE001, the filgrastim insert was amplified from its gBlock using primers F2/R2 which contain ~20 bp homologous to pBE-S. Similarly, for pAVE003, the teriparatide drug cassette insert was amplified from its gBlock using primers F2/R3. Following pBE-S PCR linearization, 1 µL DpnI (New England Biolabs, Ipswich, MA) was added per 50 µL of PCR reaction and the reaction was incubated at 37°C for 2 hours. Inserts and backbone were run on a Tris-borate-EDTA 1% agarose gel containing ethidium bromide to stain the DNA. Bands of expected sizes were excised from the gel and impurities were removed using a gel extraction kit (see above) and eluted in nuclease-free water. The resulting DNA was quantified and used in In-Fusion cloning reactions according to manufacturer protocol (incubation at 50°C for 15 minutes). In-Fusion reactions were used for transformation into *E. coli*.

#### 3.2.2 Preparation of Signal Peptide Libraries and isolation of pAVE055 and pAVE061

pAVE001 and pAVE003 were mini-prepped from single colonies of *E. coli* following confirmation *via* Sanger sequencing. pAVE001and pAVE003 were both PCR-linearized in 50 µL reactions using primers F4/R4 and the resulting PCR reactions were incubated with 1 µL DpnI at 37°C for 2 hours. Following this, reactions were run on a Tris-borate-EDTA 1% agarose gel containing ethidium bromide to stain DNA. Bands of the correct size were excised from the gel and cleaned using a gel extraction kit (see above) and eluted in nuclease-free water. The resulting DNA was quantified. A library of 173 unique signal peptides found in *Bacillus subtilis* strain 168 was inserted into this linearized backbone using In-Fusion cloning and the Signal Peptide Library DNA purchased from TAKARA Bio as a part of their *B. subtilis* Secretory Protein Expression System, according to the manufacturer’s protocols. Two µL of the resulting reactions were used directly to transform *E. coli* cells. Colony-containing plates (> 2,000 colonies) were washed and scraped into 3 mL of Luria-Bertani (LB) broth three times per plate and washes were pooled into a 50 mL falcon tube according to manufacturer’s protocol. The pooled colonies were then centrifuged, the supernatant was discarded, and the resulting cell pellets were mini-prepped to obtain the plasmid library. The resulting library was stored at -20°C for *B. subtilis* transformation and subsequent screening (see HiBiT Bioassay subsection for more detail). The best performing colonies were teriparatide colony 441 and filgrastim colony 432. Plasmids pAVE055 and pAVE061 were mini-prepped from these colonies, respectively.

#### 3.2.3 pAVE114 and pAVE118 Construction

Plasmids pAVE055 and pAVE061) were mini-prepped from colonies that exhibited the highest secretion of teriparatide and filgrastim, respectively, as evidenced from luminescent screening (see HiBiT Bioassay subsection for more detail). These plasmids were Sanger sequenced and used to clone the drug cassette and favorable signal peptide into pECE321, a suicide plasmid for ectopic integration (Guiziou et al., 2016). pECE321 was purchased from the Bacillus Genetics Stock Center (BGSC, Columbus, OH). To create pAVE114 and pAVE118, pECE321 was PCR linearized with primers F5/R5. Following a DpnI digest, this backbone was gel purified. Plasmids from the best performing colonies for each drug were both amplified using primers F6/R6 to create inserts. Inserts and backbone were electrophoresed on a Tris-borate-EDTA 1% agarose gel containing ethidium bromide to stain DNA. Bands of the correct size were excised from the gels and cleaned using a gel extraction kit (see above) and eluted in nuclease-free water. The resulting DNA was quantified and used in In-Fusion cloning reactions according to manufacturer’s protocols (incubation at 50°C for 15 minutes). A 1 µL aliquot of In-Fusion reaction was used for transformation of 50 µL of chemically competent *E. coli* cells.

#### 3.2.4 pAVE147 and pAVE148 Construction

The same plasmid used to create pAVE114 was mini-prepped and used to clone the drug cassette and favorable signal peptide into pECE321. Multiple cloning efforts were carried out using *E. coli* as an intermediate host for replication prior to mini-prepping for *B. subtilis* transformation. All genomic insertions contained deleterious mutations. One genomic insertion was confirmed to have a premature stop codon in the signal peptide region. This was used as a negative control moving forward. All other aspects of the plasmid were confirmed to be unchanged by Sanger sequencing. This control plasmid is referred to as pAVE148.

To create pAVE147, pAVE148 was PCR linearized with primers F7/R7 which overlap at the mutation site to restore the proper translation frame. Following a DpnI digest, this backbone was gel purified. This amplicon with the reversed mutation was circularized by In-Fusion® Snap Assembly according to manufacturer’s protocol. To acquire sufficient DNA for *B. subtilis* transformation (2 µg), the resulting reaction mix was diluted with nuclease-free water (4:1) and 1 µL of the diluted aliquot was used as the template in several 50 µL PCR reactions using primers 9F/9R to amplify the portion of the plasmid necessary for homologous recombination into the *B. subtilis* genome. The PCR reactions were then gel electrophoresed and DNA bands of expected sizes were removed from the gel using a gel extraction kit (see above) and eluted in nuclease-free water. The resulting DNA was quantified and 2 µg was used for transformation of 600 µL of naturally competent *B. subtilis* cells.

#### 3.2.5 *B. subtilis* Colony PCR to Confirm AE147 and AE148

*B. subtilis* cells were transformed from the DNA mentioned previously to allow for integration of the constructs present in pAVE147 and pAVE148 into the *B. subtilis* genome at the *amyE* locus. This was done to enable stable transgenic expression of teriparatide.

To confirm correct ectopic integration of final cassettes into the *B. subtilis* genome, colony PCR of strain genomic DNA was carried out. In brief, individual colonies were picked and scraped into PCR tubes containing 10 µL of nuclease nuclease-free water. Colonies were also used to inoculate selective LB media for making glycerol stocks. PCR tubes containing the cell-water suspensions were then incubated on ice for 5 minutes, then microwaved on full power for 1 minute, then chilled on ice for 30 seconds. The 1 minute microwaving and 30 second ice incubation process was then repeated for a total three times in the microwave oven. Tubes were then chilled on ice for 5 minutes. A 1 µL aliquot of microwaved cell-water suspension was then used in a 50 µL PCR reaction. PCR reactions were done according to manufacturer’s protocol for PrimeStar DNA Polymerase with the following changes: a 10-minute incubation at 95 °C was added to the beginning (before cycling). Immediately after the PCR reaction was complete, EDTA was added to a final concentration of 50 mM to prevent heat resistant DNAse degradation released from *B. subtilis* cytoplasm. Colonies that demonstrated expected sequences from colony PCR and Sanger sequencing were named strains AE147 and AE148.

### 3.3 Bacterial Transformation

#### 3.3.1 *E. coli* Transformation

A 100 µL aliquot of *E. coli* Stellar™ Competent Cells (TAKARA Bio Inc., San Jose, CA) was added to pre-chilled centrifuge tubes and chilled on ice for 10 minutes. In-Fusion Reaction Master Mix (2 µL) was then added into each tube and chilled on ice for 30 minutes. Competent cells were heat shocked on a thermal block halfway filled with water to 42°C for 45 seconds, then chilled on ice for 5 minutes. A 900 µL volume of SOC medium, pre-warmed to 37°C, was added into each tube and placed in a shaking incubator at 37°C and 250 rpm for 60 minutes. For signal peptide library creation, several 200 µL aliquots were plated on LB + Amp [100 µg/mL] selective plates. The plates were incubated at 37°C for 16 hours.

#### 3.3.2 *B. subtilis* Transformation

SPI medium, SPII medium and 100 mM EGTA (pH 7.0) were prepared according to the manufacturer’s instructions for the B. subtilis Secretory Protein Expression System (TAKARA Bio). For signal peptide library transformation into *B. subtilis*, all steps in manufacturer protocol were followed. For our inserted target genes, 600 µL plating volume yielded a sufficient number of wells-spaced colonies for screening. Cells were plated on LB + Km [10 µg/mL] selective plates.

For ectopic integration, all steps outlined in the aforementioned protocol were followed, however, after inoculating SPII medium and incubating at 37°C for 90 minutes, the cell suspension was divided in 600 µL aliquots in microcentrifuge tubes. Glycerol was added to 10% (v/v) and aliquots were flash frozen in liquid nitrogen before being stored at -80°C for later use. When ready to use in transformations, tubes were incubated at 37°C until completely thawed and 6 µL of 100 mM EGTA (pH 7.0) was added. Tubes were then shaken at 37°C, 90 - 100 rpm for 5 - 10 minutes before adding 2 µg of plasmid to be recombined into the host genome. After DNA addition, the cell suspensions were incubated at 37 °C, 90 - 100 rpm for 90 minutes. A 200 µL aliquot was then used for plating on Sp [100 µg/mL] selective LB plates overnight. Individual colonies were also streaked on Em [1 µg/mL] selective LB plates for counterselection. Colonies which grew on Sp selective plates but did not grow on Em selective plates were picked for colony PCR. Colony PCR was done to amplify the genomic DNA using primers F10/R10 and resulting amplicons were sent for Sanger sequencing using sequencing primers F11 and R11.

### 3.4 High-Throughput Bioassay for Detecting Protein Secretion

A 150 µL volume of LB + Km [10 µg/mL] was inoculated with single *B. subtilis* colonies in 96-well plates. Border wells were not inoculated as they experienced significant evaporation during growth. Three control colonies were added to each individual plate to use as a reference and to monitor ‘plate-to-plate’ variability in data. These controls were: *B. subtilis* strains harboring pAVE001 and pAVE003 (pBE-S containing HiBiT-tagged filgrastim and teriparatide cassettes with a known low-level secretion signal peptide from the aprE gene) and a *B. subtilis* strain harboring an empty pBE-S backbone (no drug or HiBiT tag inserted). Each plate also contained a blank LB control. Cultures were grown at 37 °C for 24-48 hours.

Following growth, HiBiT reagent was prepared according to manufacturer directions (Promega Corp.). Using a multichannel pipette, 25 µL of freshly made HiBiT reagent was added to new 96-well white opaque plates used to detect bioluminescence (Fisher Scientific). Corresponding wells in the growth plates were mixed well by pipetting up and down before transferring the same volume (25 µL) of culture. Following culture additions, dispensed cultures were mixed well with the HiBiT reagent by pipetting. The culture-reagent mixture was allowed to incubate at RT for no longer than 10-12 minutes before being measured on a plate luminometer using default instrument parameters.

Following the luminescent reading, luminescence values read from blank LB controls were averaged and the average value was subtracted from all other well readings to correct background luminescence. Luminescence data was then analyzed, and the brightest colonies were saved from the remainder of culture plates to make 15% (w/v) glycerol stocks stored at -80°C. All stored colonies were streaked on LB + Km [10 µg/mL] plates to isolate single colonies for miniprepping and further screening.

### 3.5 Secretion vs. Temperature

Two separate pre-cultures (5 mL LB + Sp [100 µg/mL]) were inoculated from glycerol stocks of *B. subtilis* strains AE147 and AE148 and grown for 16 hours at 28 °C. Following the 16-hour incubation, six 5 mL expression cultures were started for each of the two strains by inoculating 5 mL of Sp containing LB with 50 µL of pre-culture. These were incubated at temperatures ranging from 15-40 °C in 5 °C increments for 24 hours with shaking at 200 rpm. After growth, triplicate 50 µL aliquots were taken from each culture and absorbance readings at 600 nm (OD600) were taken. Following OD_600_ readings, triplicate 25 µL aliquots of each culture were added to white, opaque 96-well plates and reagents were added according to the manufacturer’s protocols for the Nano-Glo® HiBiT Extracellular Detection System (Promega Corp.). After a 10-minute incubation step, luminescence was read from plate wells. Triplicate blank LB readings were also conducted and the average of these readings were subtracted from all other luminescence values prior to analyses.

### 3.6 Spore Preparation and Quantification

Spores were prepared by adapting a previously published protocol (Munakata et al., 1991) as follows. A colony was inoculated into 25 mL of freshly prepared DSM and grown at 37°C and 150 rpm for 2 hours. This pre-culture was diluted 1:10 into 250 mL of prewarmed (37 °C) DSM in a 2 L flask. This larger culture was incubated for 48 hours at 37 °C and 150 rpm. The entire culture was then centrifuged for 10 minutes at 9,400 xg at 4 °C and the supernatant was carefully discarded. The resulting pellet was washed with 200 mL of cold (4 °C) sterile DI water via resuspension. The culture was then centrifuged again, and the supernatant was carefully discarded. The pellet was once again resuspended in 200 mL of fresh cold sterile DI water and the resuspended pellet was incubated overnight 4 °C. Following overnight incubation, water/spore suspension was again centrifuged at 9,400xg, supernatant was carefully discarded, and the pellet was resuspended in 200 mL cold sterile DI water. The overnight incubation at 4 °C was repeated once more. Spores were then examined microscopically with a Zeiss Axio Imager Z1 under phase contrast optics and the vast majority of suspension was observed to be free-living spores. Finally, the suspension was centrifuged once more, the supernatant was discarded, and the pellet was resuspended in 20 mL of 70% ethanol and the lids of storage tubes were wrapped with parafilm to prevent ethanol evaporation. Spore suspensions were allowed to incubate at 4 °C for 1 hour prior to plating to ensure vegetative cell death. Aliquots of each spore suspension were diluted 1-million-fold in 70% ethanol and 20 µL of this 1-million-fold dilution was plated on selective solid media, then incubated at 37 °C overnight. Resulting colonies were counted to determine the number of colony forming units (CFU)/mL of each spore suspension.

For paper blots containing *B. subtilis* endospores, ~2 cm^2^ pieces of normal printer paper were cut into small squares, folded into aluminum foil, and autoclaved. Aliquots of 10 µL of spore suspensions were blotted onto each piece of paper and allowed to dry in a sterile culture flow hood. These pieces of paper were then re-wrapped in sterile aluminum foil, labeled, and stored in 50 mL falcon tubes at room temperature for future use.

### 3.7 Pharmaceutical Peptide Production

For analysis of optimal expression temperature, a single bacterial colony was picked from each plate and suspended separately in 5 mL of streptomycin-containing LB broth and incubated overnight at 37°C. A 10 mL volume of starter culture was added to 50 mL antibiotic-containing LB in a 250 mL conical flask and incubated at 37°C for 24-48 hours. The mixture was transferred to a 50 mL Falcon tube and centrifuged at max speed for 5 minutes; supernatant was collected in a clean 50 mL Flacon tube for purification.

Expression cultures used to produce Table 1 were made by inoculating fresh media with dried spore blots and allowing the cultures to grow for 24 hours before purifying the secreted protein. Dried *Bacillus subtilis* spores prepared from strains AE068 (pBE-S_YoqHSP-HiBiT-Filgrastim) and AE147 (Δ*amyE*, +WalMSP-HiBiT-Teriparatide) were blotted on pieces of paper and stored dried at room temperature. These paper blots were used to inoculate 50 mL cultures in 250 mL conical flasks with fresh selective LB media. Cultures were grown for 24 hours at either 25 °C or 37 °C before being spun down in 50 mL Falcon tubes (9400xg, 10 minutes). The resulting supernatant was transferred to a clean 50 mL Falcon tube for purification.

**Table 1.** Maximum productivity rates observed from dried B. subtilis spores

| Strain | Drug | Growth Temperature (°C) | Productivity (ug/mL/day) |
| --- | --- | --- | --- |
| AE147 | Teriparatide | 25 | 14 |
|  |  | 37 | 11 |
| AE068 | Filgrastim | 25 | 4 |
|  |  | 37 | 5.8 |

### 3.8 Protein Purification and Quantification

An 8 mL aliquot of each supernatant was added to the same volume of standard equilibration buffer (20 mM Tris-HCl, 300 mM NaCl, antiproteases, pH 7.5). This mixture was then loaded onto a HisPur™ Ni-NTA Spin Column and the protein was purified according to the manufacturer’s instruction with the following buffer changes: Wash buffer used was 20 mM Tris-HCl, 300 mM NaCl, antiproteases, pH 7.5. Elution buffer used was 20 mM Tris-HCl, 300 mM NaCl, 250 mM Imidazole, antiproteases, pH 7.5. The flowthrough, column wash, and elution fractions were collected and analyzed by SDS-PAGE to check for sufficient purification of filgrastim and teriparatide (Fig. 9). Marker used was Novex Sharp Pre-Stained Protein Standard (Invitrogen) SDS-PAGE gel analysis showed isolated protein bands at the expected molecular weights in the elution fractions. Following His-tag purification, the eluted fractions were combined and concentrated to 2-4 mL final volume using a 5-20 mL, 3K MWCO Pierce Protein Concentrator according to manufacturer’s instruction. Imidazole was removed using a 10 mL 7K MWCO ZebaSpin Desalting Column according to manufacturer’s instruction. During desalting, a buffer exchange was conducted to transfer to protein to a 2x storage buffer (40 mM Tris-HCl, 600 mM NaCl, antiproteases, pH 7.5). All of the resulting buffer exchanged protein was then concentrated to ~75 µL final volume using 0.5 mL, 3kDa Amicon Ultra-Centrifugal Filters according to manufacturer’s instructions. Protein quantification was conducted via the Thermo Scientific Pierce™ Rapid Gold BCA Protein Assay Kit in 96-well plates using 10 µL of each purified protein solution. Following quantification, protein solutions were diluted 1:1 with glycerol for a final concentration of 50% glycerol (w/v) and 1 x storage buffer. Proteins were then stored at -20 °C.

## 3 Results

### 3.1 Secretion Peptide Screening

Both filgrastim and teriparatide were fused to a library of secretion peptides with a HiBiT reporter tag to detect secretion via luminescence. In the first round of colony screening, 277 colonies expressing filgrastim constructs and 368 colonies expressing teriparatide constructs were individually picked, grown, and screened from the resulting library. This corresponded to three 96-well plates for filgrastim colonies and four 96-well plates for teriparatide colonies (Figs. 2 and 3). Each plate contained its own set of controls including: 1) a positive control colony harboring either filgrastim or teriparatide fused to a HiBiT reporter tag and a secretion peptide from the *aprE* gene inserted in pBE-S (i.e., aprESP_Fil/Ter). 2) a negative control colony harboring empty pBE-S with no insert (i.e., Empty_BB). 3) uninoculated media for subtracting background luminescence from readings. Following a 24-hour growth period, plates were screened for luminescence, indicative of secretion. In this initial screen, colonies which had stronger luminescent signal compared to the aprESP_Fil/Ter controls were considered for further screening and sequencing.

**Figure 2.**
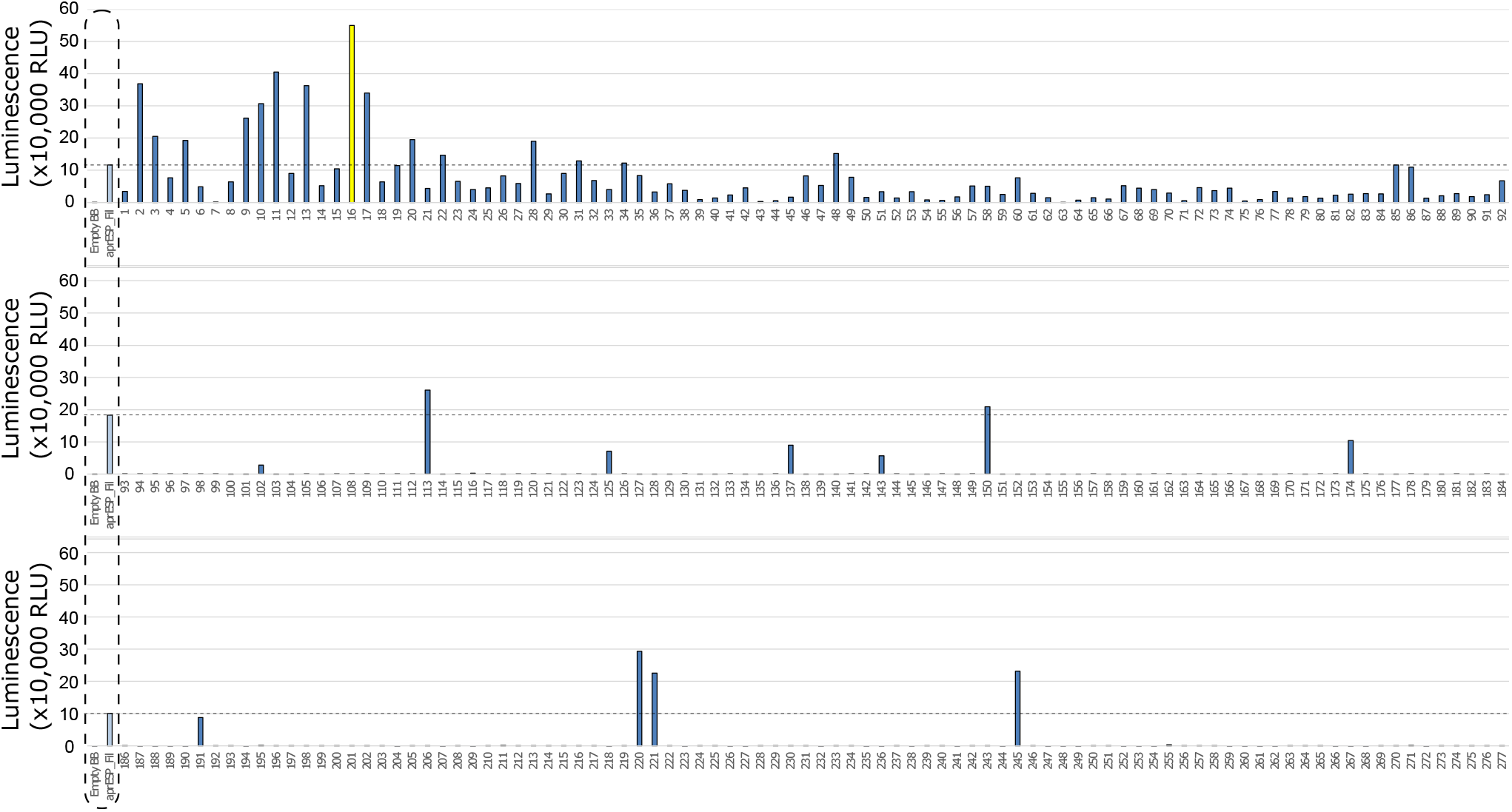
First round of colony screening of *B. subtilis* colonies secreting filgrastim fused to a HiBiT reporter and SP library. Each row shows 1 plate screened. Colonies are numbered below x-axis. Dotted box shows negative and positive controls (from l left to right). Controls were included separately in every plate. Negative control is an empty backbone control without filgrastim fusion peptide insert. Positive control are cells expressing filgrastim fusion peptide tagged with the HiBiT reporter and a secretion peptide from the *aprE* gene. Dotted line shows luminescent signal from positive control. Colonies with luminescence above this contained secretion tags with better performance compared to the positive control. Colony 16 is shown in yellow and was considered the best performing candidate to take forward for further screening and sanger sequencing.

**Figure 3.**
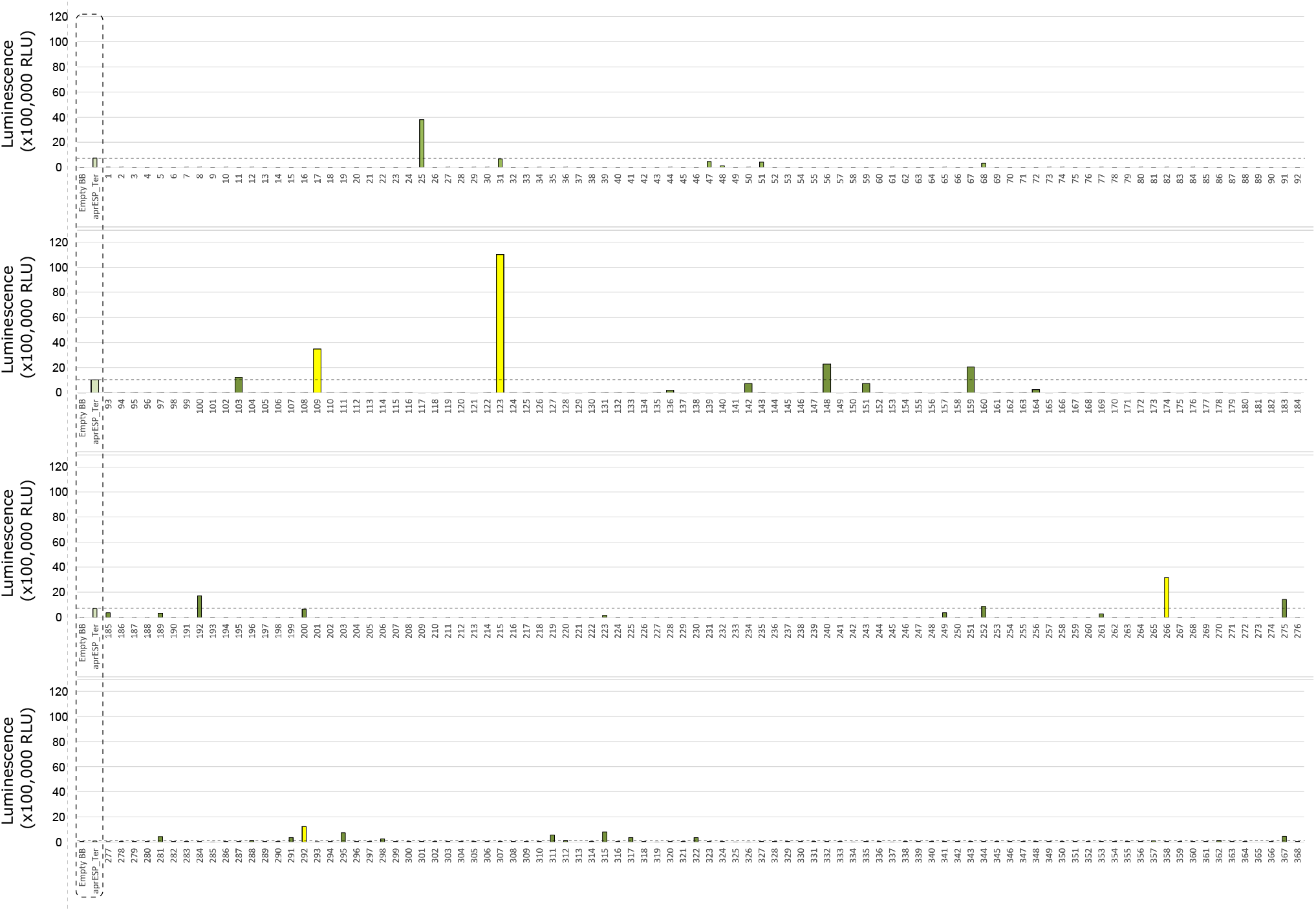
First Round of Colony screening of B. subtilis colonies secreting teriparatide fused to a HiBiT tag and one of 167 secretion peptides. Each row shows 1 plate screened. Colonies are numbered below x-axis. Dotted box shows negative and positive controls (from left to right). Controls were included separately in every plate. Negative control is an empty backbone construct without teriparatide fusion peptide insert. Positive control are cells expressing teriparatide fusion peptide tagged with the HiBiT reporter and a secretion peptide from the *aprE* gene. Dotted line shows luminescent signal from positive control. Colonies with luminescence above this level contained secretion tags with better performance compared to the positive control. Colonies 109, 123, 266, and 292 are shown in yellow and were considered the best performing candidate to take forward for further screening and sanger sequencing.

One colony stood out from the filgrastim library: colony 16 (Fig. 2). This colony gave a reading of >550,000 relative luminescence units (RLUs) (Supplement File 1). This corresponded to an over 5-fold increase in RLU signal as compared to the positive control. Due to the higher positive control variability seen in the teriparatide library plates, 4 colonies were selected for further screening (Fig. 3). Colonies 109, 123, 266, and 292 all exhibited RLU values well above the aprESP_Ter positive control (~3.5, 11, 3, and 1 million RLUs, respectively) (Supplementary File 1). These readings corresponded to an over 3, 10, 4, and 18-fold increase in RLU compared to positive controls, respectively. Sanger sequencing revealed that filgrastim colony 16 contained the secretion peptide from the *mpr* gene (UniProt: P39790). Secretion peptides from teriparatide colonies 109, 123, 266, and 292 were from genes *phrG, sacC, yncM, ypuA* (UniProt: O32295, P05656, O31803, P31847), respectively.

Following the first round of colony screening, an additional 457 filgrastim colonies and 448 teriparatide colonies (corresponding to five 96-well plates per drug) were screened in the second round (Figs. 4 and 5). Colonies selected from the previous round of screening were grown up alongside unknown colonies in the next screening round in triplicate for comparison, along with 10 replicates of previously included positive and negative controls (one per plate). Previously identified colonies were confirmed to exhibit higher luminescence compared to the aprESP_Fil/Ter positive controls and teriparatide colony 123 showed the highest average luminescence of the previous set.

**Figure 4.**
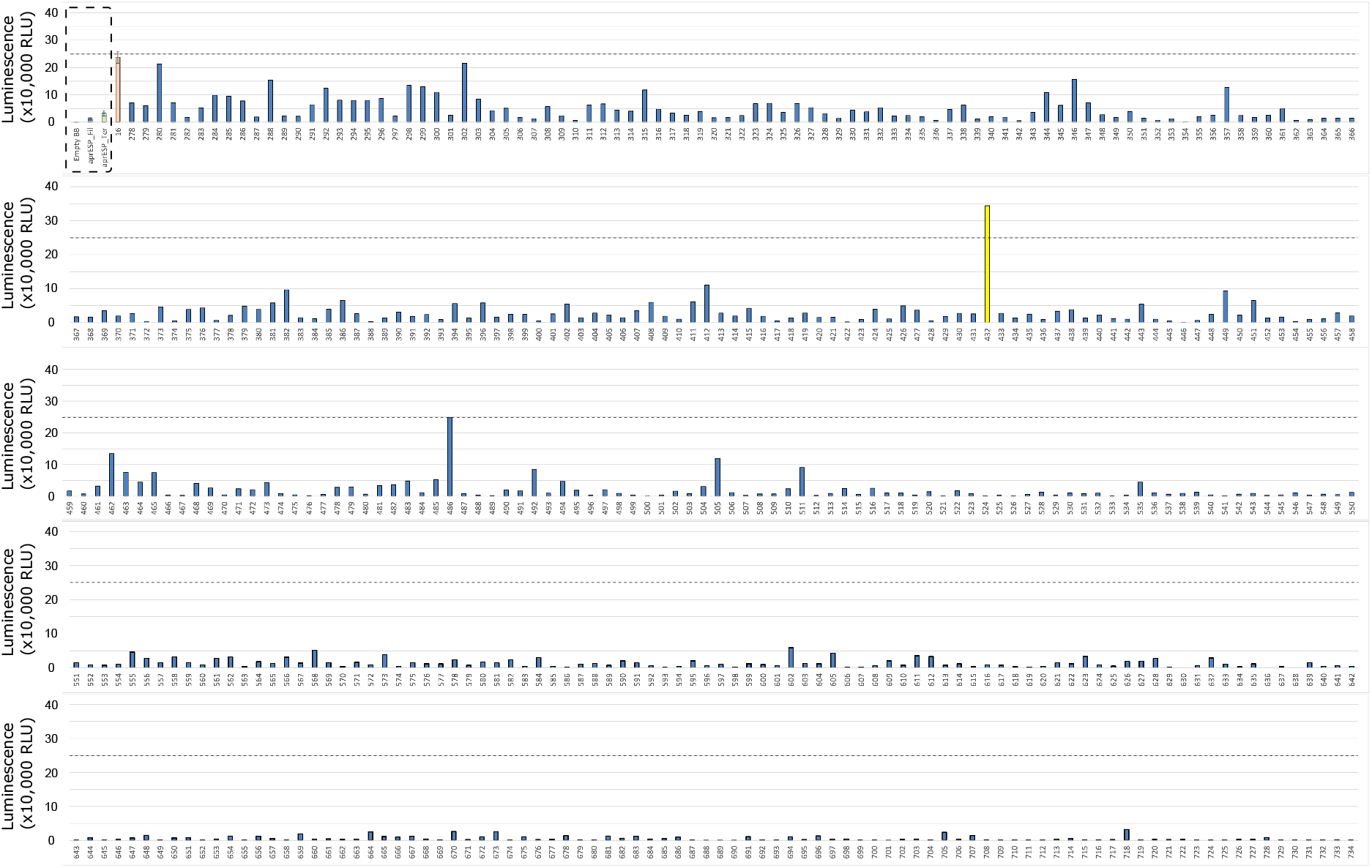
Second Round of colony screening of *B. subtilis* colonies secreting filgrastim fused to a HiBiT tag and one of 167 secretion peptides. Each row shows 1 plate screened. Colonies are numbered below x-axis. Dotted box shows negative and positive controls for filgrastim and teriparatide (from left to right). Negative control is an empty backbone construct without filgrastim fusion peptide insert. Positive controls are cells expressing either filgrastim or teriparatide fusion peptides tagged with the HiBiT reporter and a secretion peptide from the *aprE* gene. Controls were included separately in every plate and averaged. Orange bar is average luminescence reading of best-performing colony from first screen. Dotted line shows luminescent signal from most luminescent colony from previous screen (i.e., Colony 16) (see Fig. 1). Colonies with luminescence above this level contained secretion tags with better performance compared to Colony 16. Colony 432 is shown in yellow and was considered the best performing candidate to take forward for further screening and sanger sequencing. Error bars show standard error. For error bars of positive and negative controls, n = 10. For error bars of Colony 16 (orange), n = 3.

**Figure 5.**
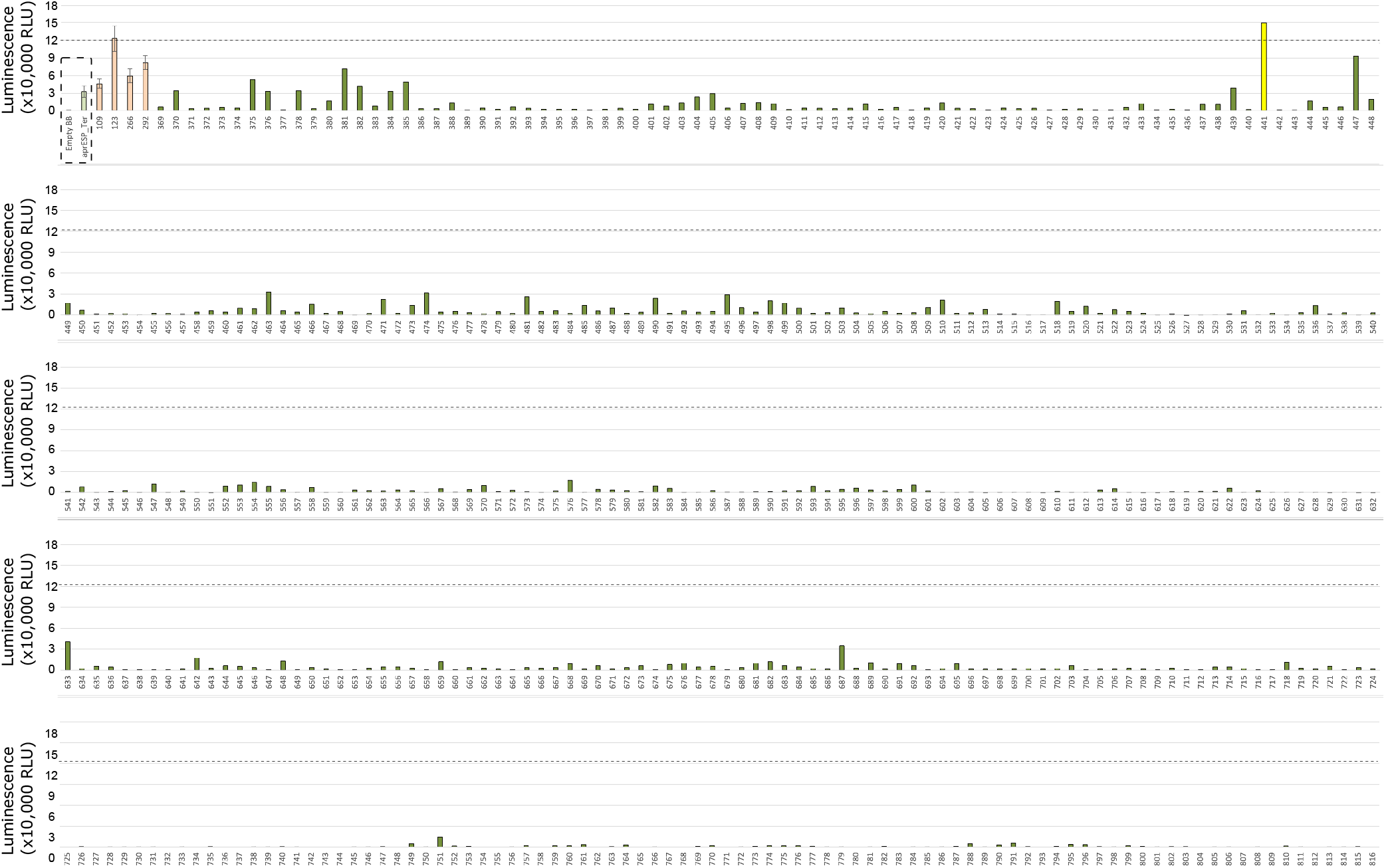
Second Round of Colony screening of *B. subtilis* colonies secreting teriparatide fused to a HiBiT tag and one of 167 secretion peptides. Each row shows 1 plate screened. Colonies are numbered below x-axis. Dotted box shows negative and positive controls for teriparatide (from left to right). Negative control is an empty backbone control without teriparatide fusion peptide insert. Positive controls are cells expressing teriparatide fusion peptides tagged with the HiBiT reporter and a secretion peptide from the *aprE* gene. Controls were included separately in every plate and averaged. Orange bars are average luminescence readings of best-performing colonies from first screen. Dotted line shows luminescent signal from most luminescent colony from previous screen (i.e., Colony 123) (see Fig. 3). Colonies with luminescence above this level contained secretion tags with better performance compared to Colony 123. Colony 441 is shown in yellow and was considered the best performing candidate to take forward for further screening and Sanger sequencing. Error bars show standard error. For error bars of positive and negative controls, n = 10. For error bars of colonies from first screen (orange), n = 3.

Filgrastim colony screening identified two colonies which exhibited higher RLU readings compared to filgrastim colony 16 from the first round of screening: colonies 432 and 486 gave RLU readings of ~340,000 and ~250,000 respectively, corresponding to a ~1.5 and ~1.1-fold increase compared to fillgrastim colony 16 (Supplementary File 2). Following sanger sequencing, both colonies were found to contain the same secretion peptide from the *yoqH* gene (UniProt: O34999) and plasmid pAVE061 was isolated from it (Fig. 6b). Teriparatide colony screening identified one colony which exhibited significantly higher RLU readings compared to teriparatide colony 123: colony 441. This colony gave an RLU reading of ~150,000 which corresponded to a ~1.2-fold increase in RLU compared to colony 123. Following sanger sequencing, colony 441 was found to contain the secretion peptide gene *walM* (UniProt: A0A6M3Z808) and plasmid pAVE055 was isolated from it (Fig. 6a).

**Figure 6.**
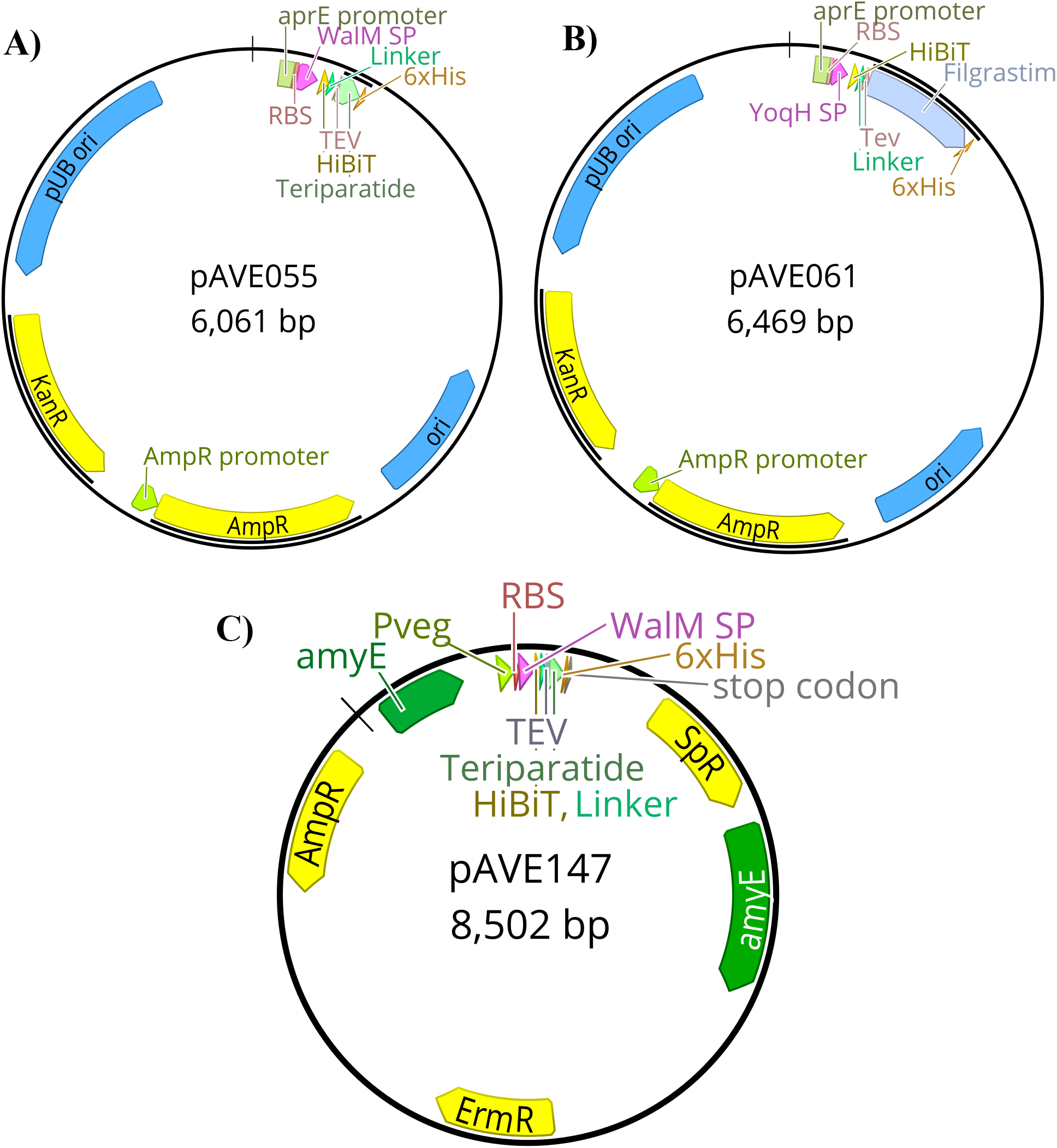
Plasmid maps of final constructs used in this study. A) pAVE055 was created by fusing the secretion peptide from the *walM* gene with a downstream HiBiT tag followed by a linker sequence and a cleavage site recognizable by TEV protease, to teriparatide tagged with a C-terminal 6x-His tag. This teriparatide fusion construct was inserted into plasmid pBE-S. B) pAVE061 was created by fusing the secretion peptide from the *yoqH* gene with a downstream HiBiT tag followed by a linker sequence and a cleavage site recognizable by TEV protease, to filgrastim tagged with a C-terminal 6x-His tag. This filgrastim fusion construct was inserted into plasmid pBE-S. C) pAVE147 was created by cloning the insert from pAVE055 into the suicide plasmid, pECE321. This plasmid has left and right-hand homology sites to the *amyE* locus of the *Bacillus subtilis* genome and functions to integrate the insert along with a spectinomycin resistance cassette for selection into this locus.

### 3.2 Genomic integration of constructs into *Bacillus subtilis*

Following identification of the best-performing secretion peptides for the filgrastim and teriparatide constructs, the constructs were inserted into the genome of *B. subtilis*. Several rounds of genomic insertion were conducted and over 40 colonies per construct were sequenced. All colonies sequenced contained frameshift mutations and premature stop codons in the secretion peptide region of the construct (data not shown). To overcome this issue, the intermediate cloning step was eliminated and replaced with PCR amplification and linearization of the construct, the product of which could directly be used for *B. subtilis* transformation (see Methods 3.2.4). This yielded a non-mutated colony for the WalMSP-Teriparatide construct (pAVE147) (Fig. 6c) which was able to be integrated into the *B. subtilis* genome successfully to produce strain AE147 (Fig. 7a). A mutant of this strain harboring a frameshift mutation in the secretion peptide region of the gene, and thus a premature stop codon, was also isolated to function as a negative control (AE148) (Fig. 7b). YoqHSP-Filgrastim construct integration showed no success using this method as dozens of colonies still possessed premature stop codons in the secretion peptide region.

**Figure 7.**
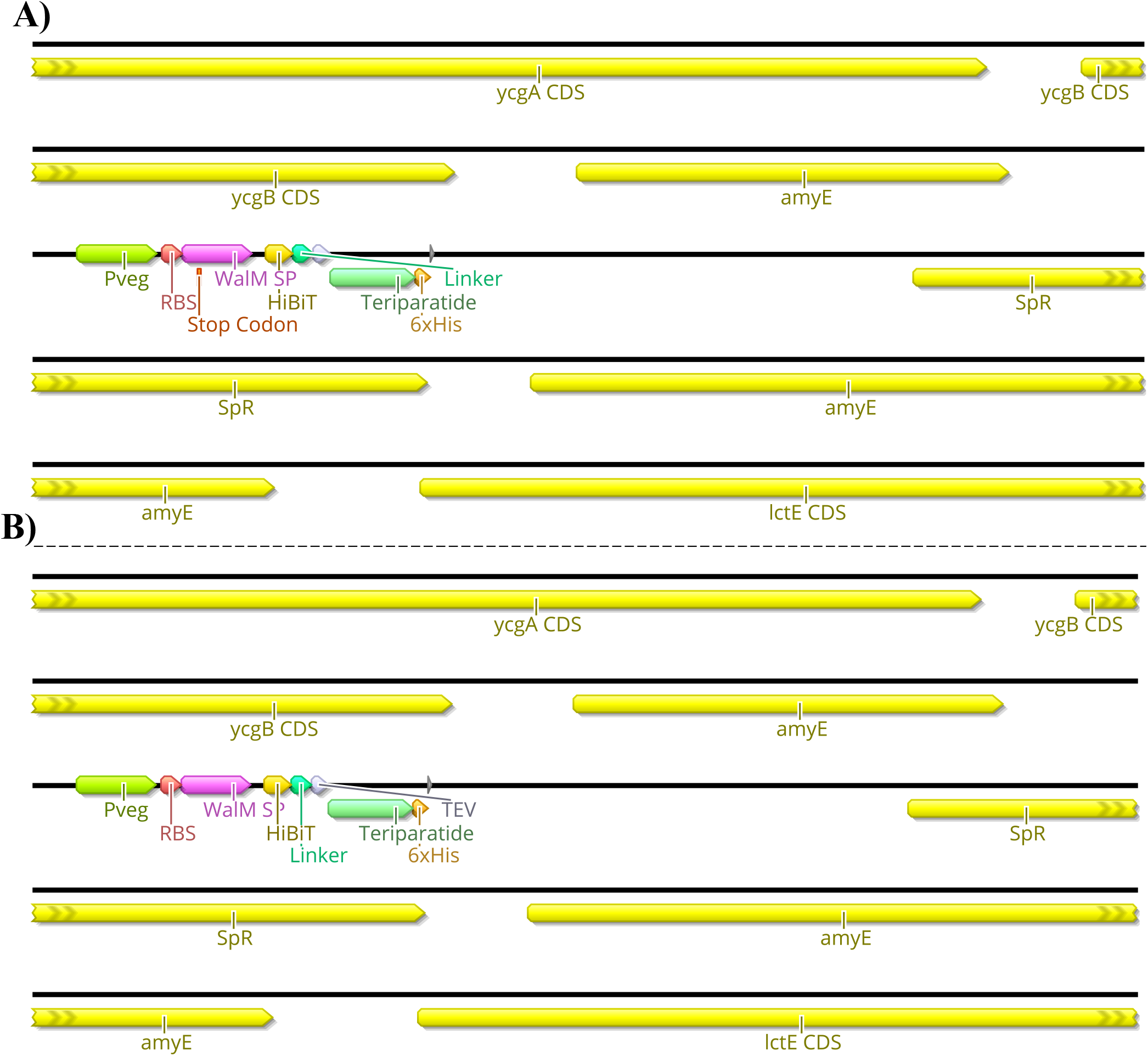
Genome Map of AE147 and AE148. This schematic shows the insertion site of the transgenic component of *Bacillus subtilis* strains AE147 and AE148. Integration was accomplished at the *amyE* locus. A) Strain AE148 consists of a truncated teriparatide secretion cassette with a single base mutation within WalM SP, resulting in a premature stop codon in the same motif. B) Strain AE147 consists of a fully functional teriparatide secretion cassette.

### 3.3 Temperature-dependent teriparatide secretion

To assess the optimal constitutive expression temperature for secretion of teriparatide, AE147 and AE148 were grown in triplicate at various temperatures and the luminescent signal was read after a 24 h incubation at these temperatures. Optical Density at the 600 nm wavelength (OD600) was also obtained after this 24 h incubation period (Fig. 8). Results showed the highest luminescent signal in cultures grown at room temperature (25 °C). Conversely, cell density was highest at 40 °C. The negative control strain showed minimal luminescence across the temperature range tested and the highest cell density was obtained when grown at 35 °C.

**Figure 8.**
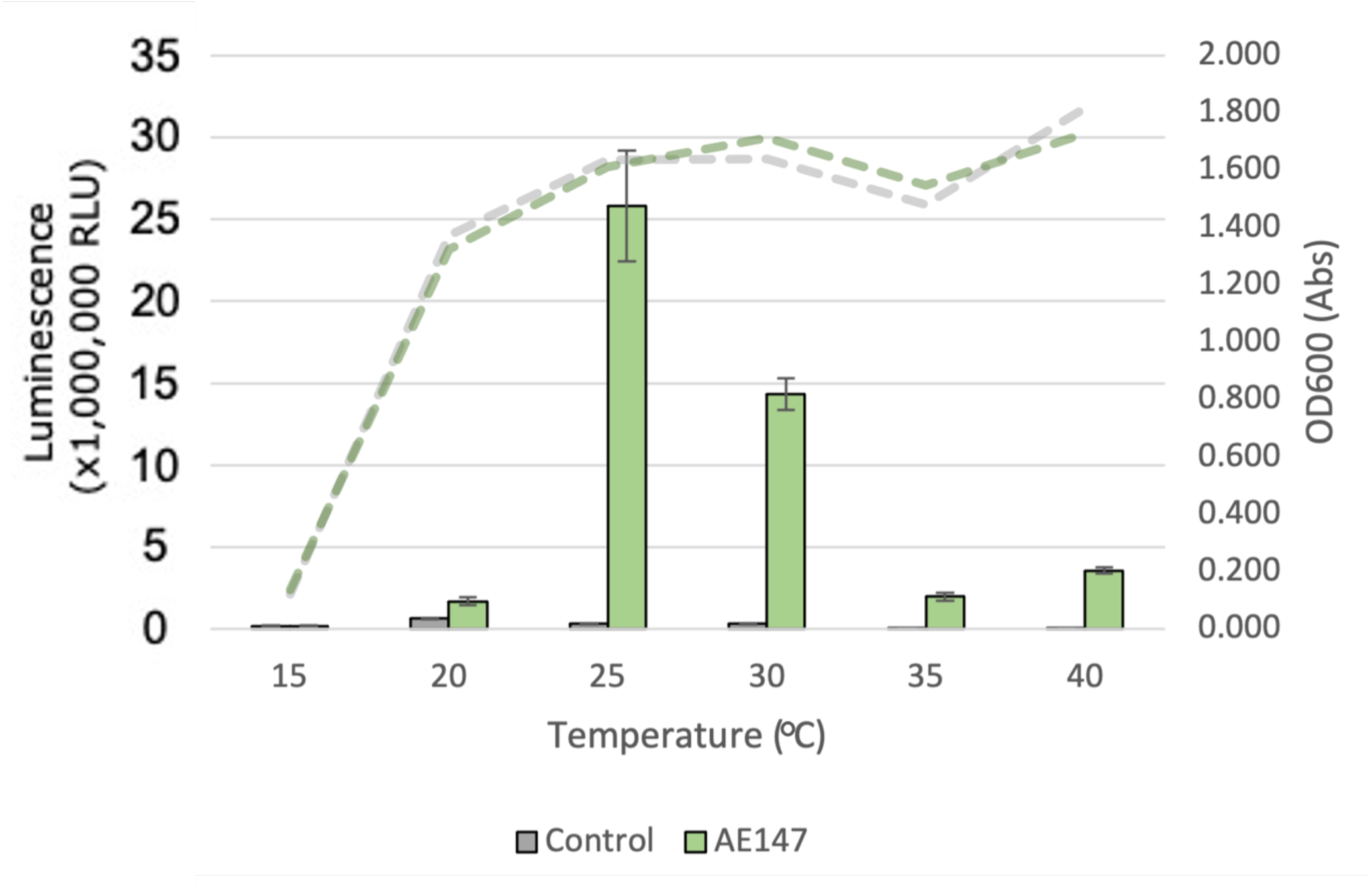
Effect of growth temperature on AE147 teriparatide secretion. Measurements are for *B. subtilis* strain AE147. This strain has a teriparatide fusion peptide inserted into the *amyE* locus. This teriparatide peptide is fused to a secretion peptide from the *walM* gene and a HiBiT reporter tag. A control strain harboring the same inserted fusion peptide but with a premature stop codon in the *walM* secretion peptide was also measured (*B. subtilis* strain AE148). Triplicate cultures were grown for 24 hours at temperatures ranging from 15 to 40 °C. Left y-axis shows luminescence values from aliquots of cultures after 24 hour incubation (bar chart). Right y-axis relates to absorbance values at OD600 taken at same time and from same cultures as luminescence values (lines chart). Error bars show standard error (n = 3).

### 3.4 Teriparatide and filgrastim purification

To test the expression stability of our strains with an eye towards implementation in our Astropharmacy, we produced spore stocks of strains AE068 and AE147. After overnight storage in 70% ethanol to rid the stocks of all vegetative cells, we blotted the spore suspension on pieces of paper and allowed the spores to air dry in the hood for 20 minutes before storing in 50 mL Falcon tubes at room temperature. Paper blots were stored for at least 7 days before being used to inoculate expression cultures. Expression cultures in Table 1 were made by inoculating fresh media with dried spore blots and allowing the cultures to grow for 24 hours before purifying the secreted protein. The flowthrough, column wash, and elution fractions were collected and analyzed by SDS-PAGE to check for sufficient purification of filgrastim and teriparatide (Fig. 9). SDS-PAGE analysis showed isolated protein bands at the expected molecular weights in the elution fractions. Sporulation and storage of our strains in desiccated conditions did not eliminate the ability of our strains to secrete filgrastim and teriparatide. The eluted fractions were then combined, and protein concentration was quantified to obtain production rates shown in Table 1.

**Figure 9.**
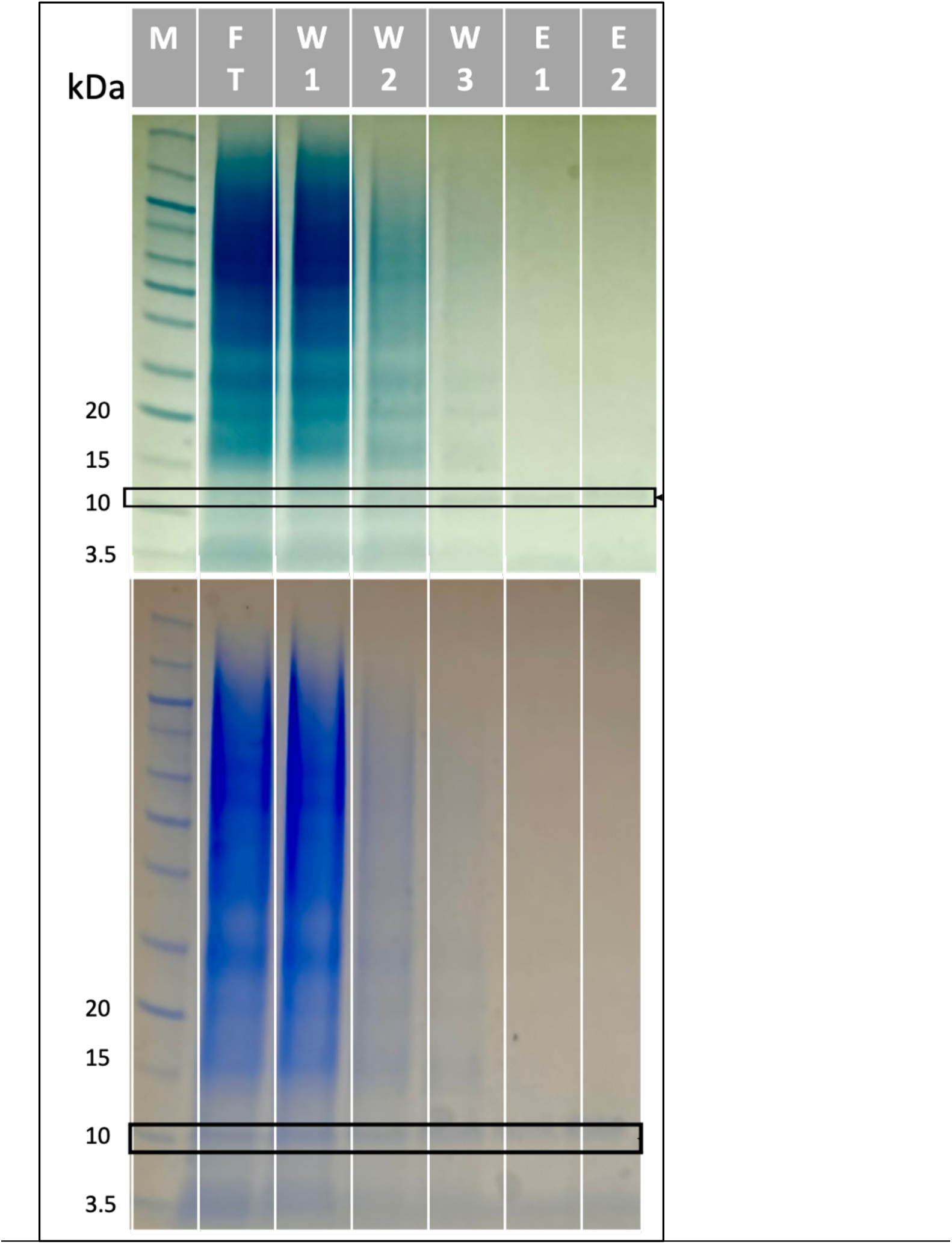
SDS-PAGE gels of filgrastim purification (Top) and teriparatide purification (Bottom). Purifications were done via His-tag spin column. M = Marker, FT = Flow Through, W = Wash, E = Elution. Protein marker used was Novex Sharp Pre-Stained Protein Standard (Invitrogen).

## 4 Discussion

By thinking of life as a technology – truly programmable, capable of self-replication and self-repair, and in some cases, dormancy (Rothschild, 2016) – we seek to develop solutions to the challenges of pharmaceutical expiration to enable more ambitious space exploration. We developed a proof of concept for a cell-based “Astropharmacy”. This platform for protein expression and secretion, where protein is manufactured according to instructions written in strands of an organism’s DNA, could produce a therapeutic dose of any protein-based biopharmaceutical within hours. Want a different drug? Use a different DNA sequence. This is the first step in developing a fully functioning Astropharmacy: the rapid on-site, on-demand production of protein-based therapeutics, and in principle, all biopharmaceuticals.

In our work, the production rate of filgrastim was less than that of teriparatide (by about 2-fold). This points to difficulties regarding filgrastim production in general which is compounded by the fact that the dose requirement for the compound is 10-fold greater than that of teriparatide. In addition to low production, we were unsuccessful in being able to move the successful filgrastim fusion-peptide into the genome of *B. subtilis*. After screening dozens of colonies for successful insertion, sequencing results showed single nucleobase frameshift mutations in the secretion peptide region of the construct. All frameshift mutations lead to premature stop codons before the CDS for filgrastim itself. This problem was thought to be a result of gene toxicity to the intermediate cloning strain being used (*E. coli*) prior to plasmid transfer to *B. subtilis*. A non-replicative cloning strategy was attempted to get around this issue. While this non-replicative cloning strategy worked for insertion of the teriparatide construct, the filgrastim construct showed the same issue in *B. subtilis* cells as it had in *E. coli* cells and, again, frameshift mutations were found in all of the dozens of *B. subtilis* colonies sequenced. This signifies that filgrastim itself is cytotoxic to *B. subtilis* cells as there seems to be strong selective pressure toward functional deletions of the recombinant filgrastim cassette when controlled with a strong genomically-localized promoter (Pveg). This cytotoxicity was alleviated when filgrastim was expressed in *Bacillus subtilis* cells via a transformed plasmid. This could be because the promoter we used for plasmid expression (P*aprE*) results in lower expression rates, allowing for cell proliferation despite filgrastim expression.

The problem of pharmaceutical toxicity to cells is a phenomenon that is seen sometimes in cell-based production systems. Not all protein-based pharmaceuticals will be amenable to recombinant expression in every strain. As a result, a large cloning effort, trying various host cells, may have to be undertaken for each new pharmaceutical needed. Conversely, cell-free expression systems do not have this problem. A great advantage of cell-free systems is that there is no need to maintain the complex physiological homeostasis of a living organism. Therefore, these systems allow for more flexibility in the type of recombinant protein it produces. However, cell-free systems have their own disadvantages. The stability of all the components in cell-free systems for recombinant protein expression is usually insufficient, particularly for the mission critical role that a life support technology, such as an Astropharmacy, would require. Whilst the PT7 expression system, which comprises just the ribosomes and 35 bacterial proteins, has shown a year-long room temperature shelf-life in lyophilized form (Pardee et al., 2014), cell extract expression systems boast shelf-life stability on the order of weeks (Gregorio et al., 2020). Even with new advances in the discovery of low-cost excipients for enhanced stability (Warfel et al., 2023), cell-free expression systems have only been shown to be stable at room temperature for up to 4-weeks, currently. In contrast, our cell-based expression system has stability at least several orders of magnitude higher given rates of *B. subtilis* survivability in their spore state (Horneck, 1993a). Future work should seek to address issues of toxicity of certain compounds while making stability of the expression system a priority.

One solution to get around the cytotoxicity of recombinant proteins would be stringent regulatory control *via* promoter optimization. While an inducible expression system would likely lead to insufficient production rates on the scale of 24 hours, weaker constitutive promoters may allow for successful genomic integration, thereby increasing stability of the expression system, and still allow for sufficient production rates through increased culture volumes. Another solution may be optimization of the expression conditions via growth at lower temperatures. Our data indicates a significant difference in luminescent signal, indicative of recombinant protein secretion, when strains were grown at lower temperatures (25 °C) (Fig. 8). When the pharmaceutical peptides were purified, yields were comparable at 25 °C as compared to the same strains grown at 37 °C (Table 1). One explanation for this may be proper peptide folding in the lower temperature condition which leads to increased solubility, secretion, and HiBiT function. However, the growth rate of *B. subtilis* at this lower temperature suffers, leading to a decrease in overall cell number and therefore a decrease in the mass of protein secreted. More work will have to be done to address this hypothesis but if supported, it signifies a large benefit to optimizing *B. subtilis* for optimal growth at lower temperatures using techniques such as adaptive laboratory evolution (ALE). ALE could not only decrease the problem of gene toxicity of recombinant proteins, but it could, at the same time, increase production rates, soluble yields, and the bioactivity of protein produced.

Further optimization of strain growth conditions, expression conditions, and promoters used could achieve higher production rates, however, these strains provide the template for further optimizations in the future. We can now use the methodology outlined to produce other drug candidates by *B. subtilis* secretion. All will have to undergo individual screening for the best performing secretion tag of a given peptide. Due to the cytotoxicity problems observed, it is likely that some pharmaceuticals will need to be localized on a plasmid, while others may be chromosomally integrated.

*Bacillus subtilis* is a promising platform for drug production for several reasons. In addition to the inherent space tolerance of its spores (Horneck, 1993a), marine strains of this same host have also been shown to have an increased capacity for secondary metabolite secretion (Ivanova et al., 1999; Mondol et al., 2013; Pandey et al., 2014; Velupillaimani and Muthaiyan, 2019). This is thought to be a result of selective evolution in the highly competitive marine environment where secretion of toxic secondary metabolites as a defense mechanism is beneficial. Marine strains of this species are also salt tolerant. Previous studies have shown that high concentrations of some salts predicted to be present in Martian brines provide UV radiation protection to *B. subtilis* cells in simulated Mars conditions (Godin et al., 2021). Aside from UV protection, brine present on Mars could be used with minimal desalination to grow *B. subtilis* cells on-site if marine strains are selected that can tolerate these higher salt concentrations. Thus, tools developed in the lab strain (strain 168) could be further transferred to marine strains with more favorable phenotypes for resistance and secretion to develop additional benefits to our engineered strains and a potential solution to potable water consumption on-site.

## 5 Conclusion

We have completed a proof of concept for the first step in our Astropharmacy: the production and secretion of peptide biologics using *B. subtilis* as a production platform. The pharmaceutical peptides chosen, teriparatide and filgrastim, are important for astronaut health. This effort represents the first time that peptide pharmaceuticals have been recombinantly produced in a space-tolerant microbe. Using high-throughput screening we identified the two secretion peptides that were most useful for teriparatide (*walM* gene) and filgrastim (*yoqH* gene) secretion. The resulting teriparatide-producing strain (AE147) produced 1 dose-equivalent (20 µg) of teriparatide in 24 hours from as little as 1.5 mL of cell culture. The filgrastim-producing strain (AE068) was able to produce 1-dose equivalent (300 µg) of filgrastim in 24 hours from 52 mL of cell culture. We showed that these production rates were unaffected by sporulation of the host cells, and subsequent storage in a desiccated state, indicating high stability and reliability in *B. subtilis*. Finally, we identified several considerations, previously unknown for recombinant expression of these pharmaceutical peptides in *Bacillus subtilis* that should be explored in future development of a cell-based Astropharmacy.

Further optimization of strain growth conditions, expression conditions, and promoters could enable higher production rates to be achieved. These strains provide templates that can be further optimized in follow-on work. Successful demonstration of *B. subtilis* as a peptide pharmaceutical production platform paves the way for the development of novel purification technology and drug delivery devices so that the vision of an Astropharmacy can be fully realized.

## 7 Author Contributions

The Author Contributions section is mandatory for all articles, including articles by sole authors. If an appropriate statement is not provided on submission, a standard one will be inserted during the production process. The Author Contributions statement must describe the contributions of individual authors referred to by their initials and, in doing so, all authors agree to be accountable for the content of the work. Please see <u>here</u> for full authorship criteria.

## 9 Acknowledgments

This is a short text to acknowledge the contributions of specific colleagues, institutions, or agencies that aided the efforts of the authors.

